# Pupillary oscillations accompany hand movement planning in naturalistic scenes

**DOI:** 10.64898/2025.12.23.696178

**Authors:** Sebastian Wiedenski, Marita Metzler, Christian Klaes, Artur Pilacinski

## Abstract

Pupil diameter changes may be a useful signal source for noninvasive studies of motion planning. However, the pupil is also sensitive to luminance changes, making it difficult to study pupillary signals under natural conditions. Here we used pupillary hippus, a luminance-insensitive measure of high-frequency pupil dynamics while human participants planned naturalistic reaches in virtual reality. We found that hippus is significantly increased during planning compared to both rest and movement execution epoch. This finding shows pupillary dynamics change during motion planning reflecting preparatory processes. Most importantly, we demonstrate the potential of a new pupillometric method to study motion planning in natural conditions.

## Introduction

Hand and eye activity is inherently related: when we interact with the environment, our eyes usually guide these interactions by selecting and evaluating interaction objects and guiding the upcoming manual actions [1]. Here, we show that pupillary dynamics consistently change during hand movement planning, indicating the increased cognitive load during movement planning.

Movement planning is the set of neural processes through which the central nervous system specifies, organises, and temporally structures motor commands prior to execution [2]. It entails identifying an intended action, encoding its spatial and temporal parameters, integrating sensory predictions, and preparing effector-specific motor programs. As such, we will use the term movement preparation and movement planning interchangeably. This process is tightly coupled with higher-order cognitive functions, including selective attention, working memory, perceptual decision making, and executive control, which collectively support the evaluation of environmental demands, the selection and inhibition of action alternatives, and the dynamic updating of motor plans in response to internal and external feedback. Movement planning is therefore a cognitively demanding task and the cognitive aspects of motor planning are usually studied through evaluating behaviour or brain activity. Yet, to our knowledge, very few studies covered pupillary activity reflecting movement planning compared to other behavioural or physiological measures. This is surprising as pupillary activity is known to reflect at least some cognitive processes involved in movement planning. From the available literature, pupil changes have been demonstrated to reflect saccade planning [3], hand movement distance [4], precision [5], [6], adaptation [7] or general arousal during the movement planning stage [8]. One difficulty, however, in studying pupillary activity during motion planning is that pupil dilation (the usual measure of pupil activity) may reflect many non-cognitive factors, including environmental luminance which is a common confound in reaching studies especially in naturalistic conditions in which lighting is dynamic and leads to uncontrolled pupil diameter changes. This confound is mitigated in traditional experimental studies in pupillometry by using controlled luminance but limits the usefulness of pupillometry in naturalistic lighting conditions.

This limitation can be remedied by approaches robust to pupil responses to environmental lighting. While under dynamic lighting conditions pupil size is modulated by luminance, resulting in slow changes to the pupil diameter [9], under stable lighting conditions rapid pupillary oscillations (hippus) were demonstrated to reflect cognitive input [10], [11]. Through careful decomposition of pupillary data into these slow and rapid changes in pupil diameter, it is possible to distinguish the pupillary changes due to cognitive load even in naturalistic lighting conditions, circumventing limitations of the classical approaches to pupillometry. Two most popular algorithms used for this are the index of cognitive activity (ICA, see: [10]) and index of pupillary activity/low-high index of pupillary activity (IPA/LHIPA, see: [11]). These methods utilise repeated wavelet transformation to decompose the signal into individual components for further analyses [12]. Here we leveraged these approaches to investigate pupillary activity during movement planning in naturalistic conditions. We specifically decided to test the LHIPA usability for the differentiation of movement planning from movement execution and rest in the context of a neuroprosthetic system and in a virtual environment with a head-mounted display and integrated eye tracker. We compared both pupillary dilation and LHIPA to assess slow and rapid pupillary changes during planning of three types of naturalistic reach and grasping movements.

## Materials and methods

### Subjects

Subjects *(N = 27, 14 male, 13 female, age range 20 to 40)* were recruited on a convenience basis.

All subjects were right-handed, had normal to corrected-to normal vision and suffered from no known psychiatric conditions or other conditions that could impair the measurements or cause discomfort and or harm to the subject. Subjects were asked to fill out a consent form and a minimal survey concerning the quality and length of their sleep, prior experience with VR, tendency for VR-induced motion sickness, as well as a self-assessment of current wellbeing, mood and drug consumption, both prescribed and recreational. All survey data was anonymised, stored in a GDPR-conform location and used exclusively for contextualising possible events. Lastly, subjects were asked not to apply make-up before the measurement.

All study procedures were accepted by the Ethical Committee of the Medical Faculty at Ruhr University Bochum.

### Computer setup

The computational load was divided onto two separate machines for the Unity Game Engine running the VR environment and data collection, respectively. They were connected via a 10 Gbit LAN network (Refer to the addendum for specifications).

Data transfer and synchronisation was done using the LabStreamingLayer (LSL) protocol.

### VR development, software and hardware

For the development of the VR environment, Unity Game Engine [13] LTS 2020.3.9f1 was used in combination with mostly free assets from the Unity store. A complete list of assets and plugins is available in addendum A.6. The physics interactions were implemented using the paid asset “Hurricane VR - Physics Interaction Toolkit” [14].

SteamVR was used for the communication between Unity and the VR goggles, as well as the room configuration. The Tobii Pro SDK was integrated into the project to collect eye data from the VR goggles in Unity. The LSL API was used for streaming measured data via the local network.

The HTC Vive Pro Eye head-mounted display (HMD) was used as VR goggles in this study due to their well working integrated eye tracker. The Vive Pro Eye has a display resolution of 1440 x 1660 pixels per eye, a refresh rate of 90 Hz (which was later interpolated to 100 Hz) and an effective field-of-view of 94 degrees.

For the controller, we opted for Valve Index Controllers due to their lightweight constructions and more intuitive interaction mechanics compared to the standard “Vive Wand” controller. Only the right hand was used, as such only one controller was used in the study.

### Eye sensor calibration

The integrated Vive eye sensor calibration task was performed both before the training phase and before the measurement process to ensure visual comfort in VR and optimal measurements during the experiment. Although subjects were permitted to use glasses, none reported the need after the configuration.

### Experimental protocol

Prior to the execution of the experiment, all subjects underwent a training session. This session included adjustments to the table and chair positions, 18 trial tasks, six of each type and a free roam period to get accustomed to movement in VR. The training session was also used to equip, test and configure the eye tracker of the VR headset as well as the LSL streams.

Instructions for the subject were given in written prompts using a virtual display in front of them. Verbal instructions were given exclusively if strictly necessary. Time, trial and cause were documented for each instance of interruption. The subject was instructed to place their hand in a semi-translucent green box and relax their arm with the aim to create a resting state, activating whenever the subject entered and left the box. The trial was deemed invalid if the subject left the box prior to the “go” signal.

The first instruction of every trial was the resting position of the hand, followed by a short waiting period and the new instruction (see Figure 2 A). Afterwards, the task object or objects appeared. The subject then had to wait for a fixed preparation duration of two seconds. Subsequently, the instruction changed to “go” only after which the subject was allowed to perform the task. Should the trial be completed successfully, a “Well done!” message appeared. If not, no message appeared, or in the case of the centre-out task, the selected item appeared red, instead of green. In this case, the subject was instructed to reposition their hand into the green box and repeat the task until successful or skip the task using the “skip” button on their right. Movement types comprised three daily actions: The wrist-rotation task, the drinking task and the center-out task (see Figure 2 B). For each task type existed slight variations. For the wrist-rotation task (Figure 2 B, 1), the subject had to grab a lever in-front of them and rotate their wrist by 45 degrees, either in clockwise, or counter-clockwise direction. For the drinking task (Figure 2 B, 2) the subject had to, depending on the variation, extend their arm and grasp either a bottle or a mug and lift it to eye-level. For the center-out task (Figure 2 B, 3) the subject had to select one of three objects which were, depending on the variation, either to their left, their center or their right by extending their arm and touching the object. For the purpose of pupillometry analyses all sub-variations of each task were analysed together.

### Data quality check

All data was compared with the notes made during the recording process to identify invalid segments due to erratic movement, loss of concentration, technical difficulties or similar issues that may have or have affected the signal quality. Trials with severe issues were removed, so were trials that showed insufficient signal quality. Subjects with over 1/3^rd^ excluded trials were excluded as a whole.

Both factors resulted in the final count of 27 subjects.

### Data preprocessing

Each data stream was separated into individual trials by comparing the data stream time stamps with the time stamps of the task markers. The resulting data streams were then separated into preparation and execution phases with constant sample lengths depending on the refresh rate of the respectable sensor. Each trial consisted of a preparation phase and an execution phase. Intermediary resting phases of flexible length but constituting at least two seconds were present between trials. Both preparation and execution phases were two seconds long each. Each preparation phase began with the appearance of the task instruction and ended with the instruction to initiate movement, which coincidentally started the execution phase. This phase ended with the finalisation of the task and a congratulatory message, followed by the resting phase. Each message to the subject was a marker which allowed for identification and separation of epochs in the data preparation. A schematic depiction of the task structure is presented in figure 1.

**Figure 1:**
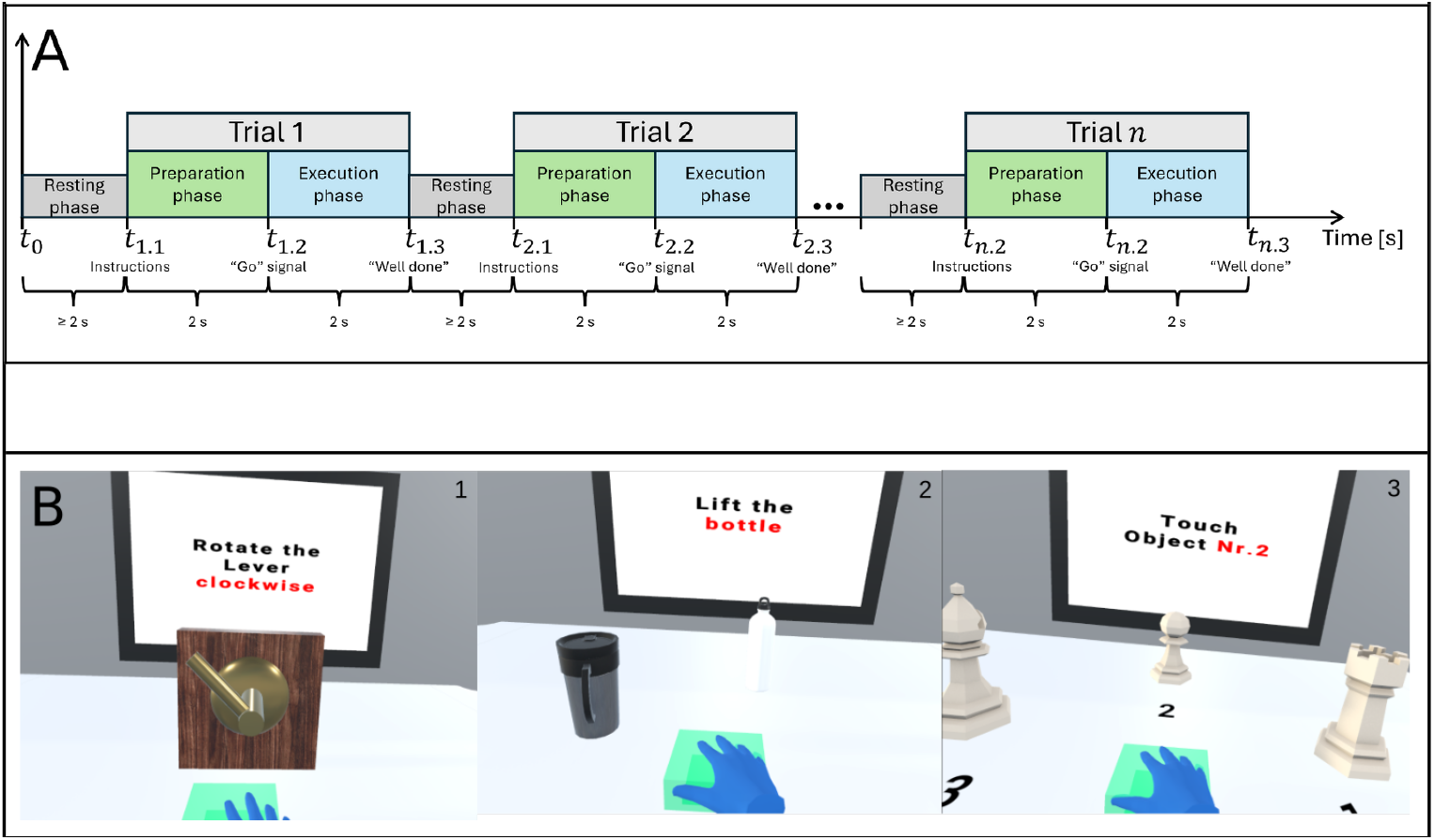
Schematic depiction of the experimental structure. Panel A: Temporal trial and epoch structure. Each trial consists of a preparation phase followed by an execution phase and an intermediary resting phase. Panel B: Task type variations as seen by the subject. 1: Wrist rotation task, clockwise-variation. 2: Drinking task, bottle-variation. 3:Center-out task, central-variation.

### Blink and invalid data removal

Based on instructions from Duchowski et al. [11], [15], we removed blinks and invalid eye signals from the raw pupil data. Invalid eye signals were caused among others by excessive gaze direction or pre-blink captures. This removal was performed in python using custom code.

Alternating from Duchowski’s method, we utilised the eye validity parameter, which illegitimises data points based on sensor detection of the pupil. This value applies a grading to the pupil data. We removed all samples with a value indicating less than perfect conditions. Adhering to that, we removed all samples including and around each blink and otherwise invalid section until the signal quality recovered. We also removed one additional sample before and after any invalid section to account for possible insufficiencies of the sensor.

### IPA and LHIPA calculation

Due to the open sourced nature of both methods, the necessary documentation and code were freely available in the original publications [11], [15]. Based on the descriptions of the papers and the similarities of our setup to the published one, we adopted the stock settings of both methods, namely the wavelet type of “symlet 16” at level two. We merely adapted the given source code to our code structure and naming convention.

### Averaging and baseline calculation

To generate a control value for both preparation and execution phase, a global baseline was calculated. For each subject, all values across the signal stream were “flattened”, meaning aligned in a row. A global median was then calculated across all trials. This median was used as the baseline for both phases and acted as the control value. In a similar manner, the medians for the preparation and execution phases were calculated by flattening all epochs of the respective phase and calculating the median per subject. These three values, the median of the preparation phase, execution phase and baseline were then used in the statistical analyses and planned as features.

### Statistical analysis

Each modality was first reviewed using Friedman’s repeated measures ANOVA to determine whether a significant difference in variance existed between any of the measured phases, followed by a Shapiro-Wilk (sw) normality test for each phase combination. Subsequently, a paired-samples T-Test was performed to identify differences between individual phases with the Student’s t for normally distributed and Wilcoxon W for not normally distributed data. Effect strength was measured by calculating Cohen’s d. Statistical analyses were performed with the open-source software jamovi [16].

## Results

### Pupil diameter

Pupil diameter results indicated a significant difference in variance among epochs, *(X* ^2^*(2) = 18*.*6, p* < .*001)*. We found significant differences between the baseline and preparation *(mean difference = 0*.*0244, T (30) = 430, p* < .*001, sw = 0*.*001, d = +0*.*545)*, baseline and execution *(mean difference = -0*.*0296, T (30) = 72, p* < .*001, sw = 0*.*005, d = -0*.*544)* and between preparation and execution *(mean difference = -0*.*0533, T (30) = 68, p* < .*001, sw = 0*.*003, d = -0*.*546)*.

Interestingly, the median of the preparation phase *(M = 2*.*77)* was smaller than both the baseline *(M = 2*.*81)* and execution phase *(M = 2*.*86)* based on the t-test results and as seen in figure 2.

**Figure 2:**
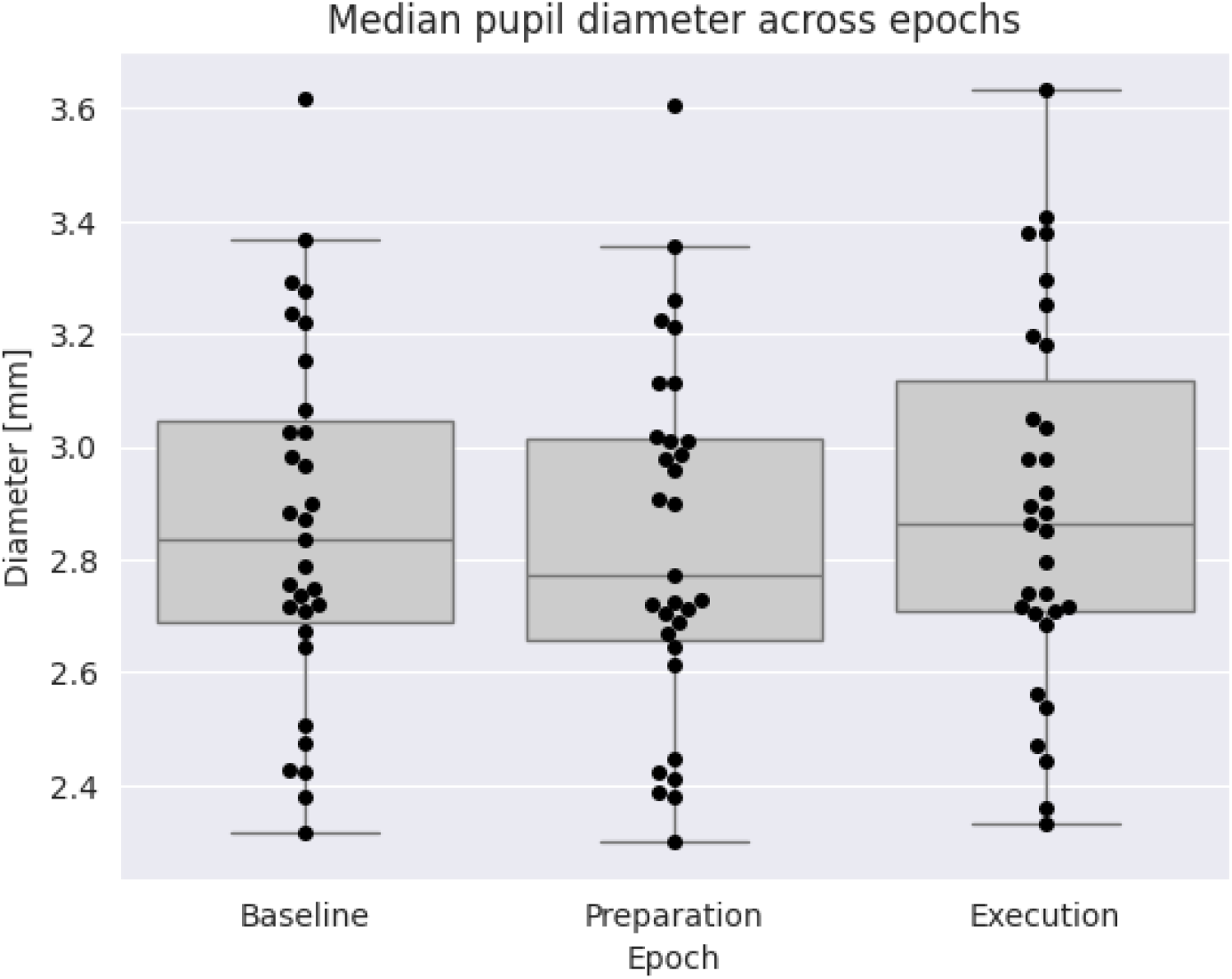
Boxplots of the pupil diameter medians during the two epochs and the global baseline. Data points in black represent the median of an individual subject. Error bars denote upper and lower quartiles, boxes denote interquartile ranges, horizontal bars denote medians.

### Low/High Index of pupillary activity (LHIPA)

Epochs of the LHIPA study exhibited a significant difference in variance *(X* ^2^*(2) = 32*.*6, p* <.*001)*. LHIPA results showed significant differences between baseline and preparation *(mean difference = 0*.*179, T (30) = 244*.*00, p* < .*001, sw* < .*001, d = +0*.*749)*, baseline and execution *(mean difference = -0*.*129, T (30) = 3*.*00, p* < .*001, sw* < .*001, d = -0*.*69)* as well as preparation and execution *(mean difference = -0*.*25, T (30) = -6*.*16, p* < .*001, sw = 0*.*077, d = -1*.*107)*.

T-test results showed that the median of the preparation phase (*M = 7*.*29*) was lower than the medians of the baseline (*M = 7*.*42*) and execution phase (*M = 7*.*55*), which can also be seen in figure 3.

**Figure 3:**
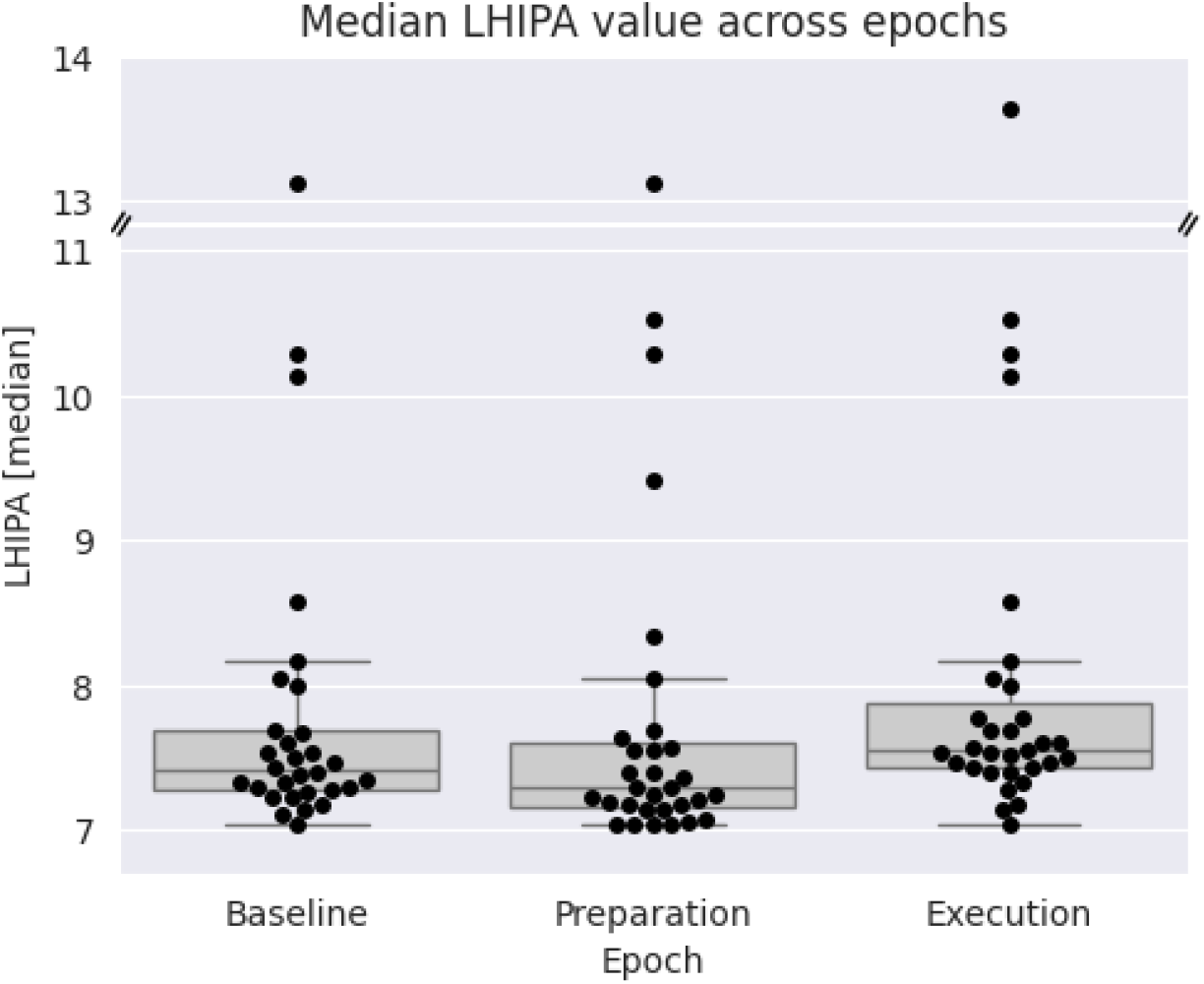
Boxplots of the LHIPA medians during the two epochs and the global baseline. Data points in black represent the median of an individual subject. Error bars denote upper and lower quartiles, boxes denote interquartile ranges, horizontal bars denote medians.

### Movement types

ANOVA analyses showed significant differences in variance in the epochs of all three tested task types, namely wrist rotation *(X* ^2^*(2) = 19*.*1, p* <.*001)*, drinking task *(X* ^2^*(2) = 23*.*0, p* <.*001) and* center-out *(X* ^2^*(2) = 22*.*2, p* <.*001)* which can be observed in figure 4.

**Figure 4:**
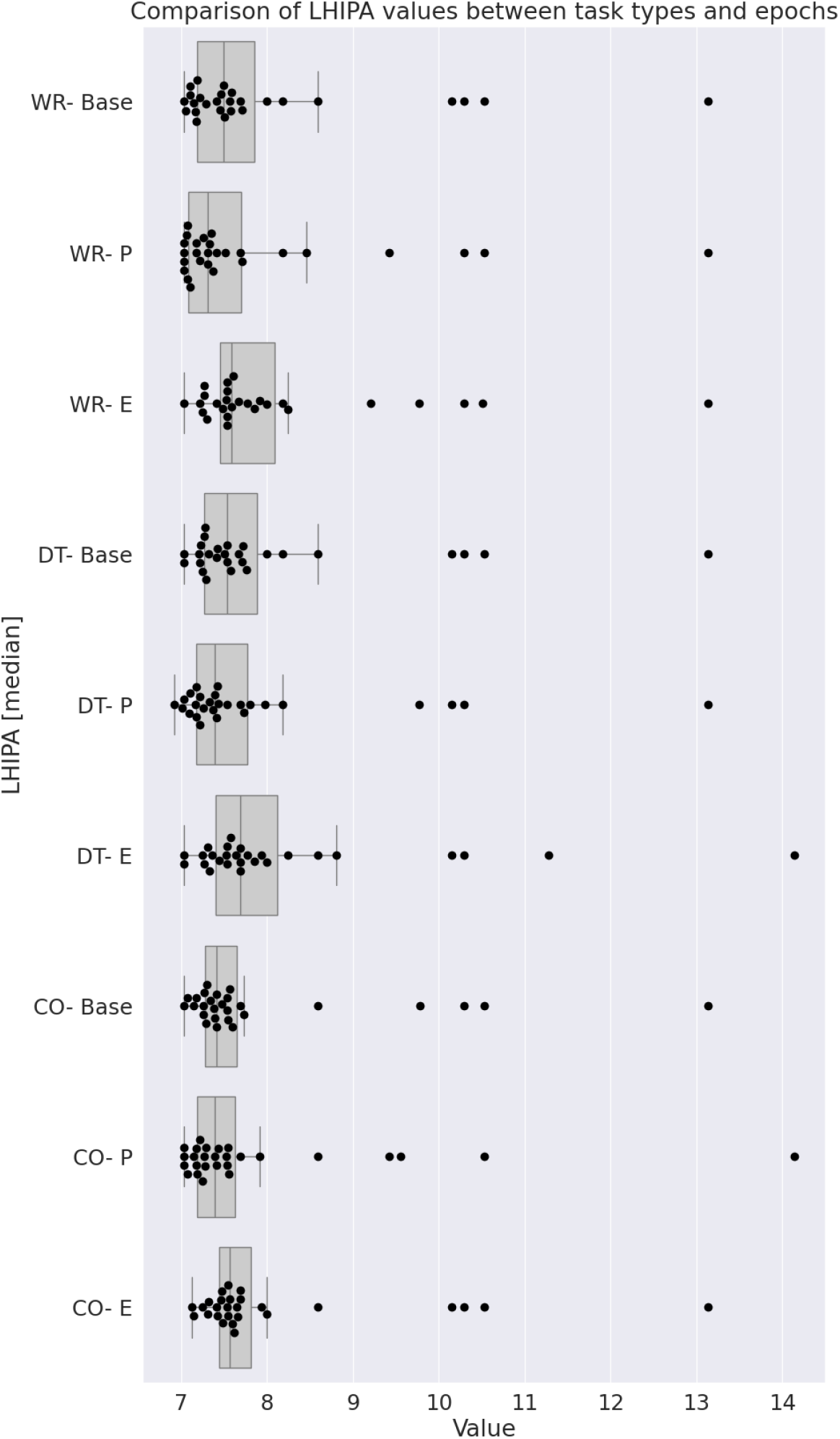
Boxplots of the LHIPA median of each trail type and epoch compared. Abbreviations: CO: Center-out task, DT: Drinking task, WR: Wrist rotation task, Base: Global baseline, E: Execution epoch, P: preparation epoch. Data points in black represent the median of an individual subject. Error bars denote upper and lower quartiles, boxes denote interquartile ranges, vertical bars denote medians.

T-test results for the LHIPA measurements further showed significant differences between all three epochs for each task type. The results showed significant differences during the wrist rotation task between baseline and preparation *(mean difference = 0*.*177, T (30) = 157*.*00, p = 0*.*002, sw* < .*001, d = +0*.*632)*, baseline and execution *(mean difference = 0*.*225, T (30) = 27*.*00, p = 0*.*004, sw = 0*.*001, d = -0*.*524)* and between preparation and execution *(mean difference = 0*.*273, T (30) = -3*.*90, p* < .*001, sw > 0*.*05, d = -0*.*750)*.

LHIPA also showed significant differences during the drinking task between baseline and preparation, *(mean difference = 0*.*176, T (30) = 173*.*00, p = 0*.*002, sw = 0*.*004, d = +0*.*661)*, baseline and execution *(mean difference = 0*.*250, T (30) = 6*.*00, p* < .*001, sw* < .*001, d = -0*.*693)* and between preparation and execution *(mean difference = 0*.*366, T (30) = 13*.*00, p* < .*001, sw = 0*.*019, d = -0*.*905)*.

As with the previous two types, LHIPA results further showed significant differences between all three epochs of the center-out task, specifically between baseline and preparation *(mean difference = 0*.*0946, T (30) = 134*.*00, p = 0*.*037, sw* < .*001, d = +0*.*182)*, baseline and execution *(mean difference = 0*.*1250, T (30) = 15*.*00, p* < .*001, sw* < .*001, d = -0*.*730)* and between preparation and execution *(mean difference = 0*.*2480, T (30) = 38*.*00, p = 0*.*004, sw* < .*001, d = +0*.*449)*.

ANOVA analyses of the LHIPA data did not however show significant differences between the three task types.

## Discussion

We investigated pupil dynamics during movement planning. We compared pupillary hippus measures with a traditional pupillometric approach and showed that pupillary hippus is increased during motion planning, or preparation, compared to baseline and motion execution. This was accompanied by decrease in pupillary diameter during movement planning compared to other epochs. These findings demonstrate that movement planning is accompanied by increased cognitive load and this is independent of general pupillary dilation. In addition, we show the feasibility of using hippus as a robust measure of cognitive processes in naturalistic motor planning tasks.

The observed pupillary changes during the interval preceding movement likely reflect an increase in cognitive load associated with preparatory processes such as action selection, specification of kinematic parameters, and anticipatory sensory prediction. This pattern is consistent with interpretations that preparatory pupil changes index the engagement of central executive resources and neuromodulatory systems that bias sensorimotor circuits toward imminent action [9], [17], [18]. Pupillary hippus, the spontaneous oscillation of pupil size, appears to arise from rhythmic fluctuations in the balance between parasympathetic and sympathetic input [19]. Evidence suggests that brainstem autonomic nuclei and neuromodulatory centers, including the locus coeruleus and midbrain structures, contribute to the amplitude and frequency of hip0pus [20]. Hippus is typically reduced during periods of sustained attention and is altered by manipulations that influence cortical excitability or autonomic tone, suggesting that it may reflect slow fluctuations in arousal systems that periodically bias motor readiness [8], [9], [21], [22].

For movement planning, the same neuromodulatory circuits implicated in pupil control interact directly with cortical and subcortical motor-planning networks. *Locus coeruleus* projections modulate prefrontal, parietal, and premotor areas by altering response gain, signal-to-noise ratios, and evidence-integration windows, all of which affect action selection and preparation [23], [24], [25]. Additionally, midbrain structures such as the superior colliculus and basal forebrain cholinergic systems influence both attentional orienting and motor preparatory activity, enabling coordinated shifts in pupil dynamics and action readiness [17], [21], [26] . Preparatory increases in pupil size or changes in hippus characteristics thus likely reflect coordinated neuromodulatory adjustments that facilitate action selection and impending motor commitment.

Pupil signals are shaped by many factors, including luminance, blink and saccade behaviour, and baseline autonomic state. Luminance introduces slow fluctuations that can confound trial-level interpretations. In movement planning studies, careful control of illumination and concurrent acquisition of behavioural or neural measures can improve the sensitivity of inferences about pupil dynamics during planning at the expense of naturalistic context of the movement involved. However, with the use of methods such as LHIPA pupillometry can successfully dissociate tonic arousal from phasic planning-related dilations and provide complementary information to neural or behavioural indices of movement preparation in both laboratory and naturalistic settings [10], [11], [15].

### Limitations

We acknowledge several limitations of the current study. First, the hippus measure did not appear sufficiently sensitive to detect differences between the movement types examined. It remains unclear whether this reflects unspecific pupillary dynamics across movement categories or instead a limitation of the eyetracking system used. Future work employing eyetracking devices with higher sensitivity and protocols specifically optimised to capture subtle differences between action types will be necessary to resolve this issue. Second, the pattern of mean pupillary responses observed here diverged from findings reported in prior research on motion planning in that in our case pupil diameter decreased prior to movement [3]. One possible explanation is that pupillary correlates of hand-movement planning may be highly context-dependent, influenced by factors such as task structure, predictability, or motor demands. This may likewise lead to distinct involvement of neural circuitry involved in regulating different types of pupillary responses. Another explanation can prove our main point of this study: pupil diameter changes are difficult to reliably capture in naturalistic conditions which may lead to spurious effects. However, given the scarcity of studies directly examining pupillary signatures of hand-movement planning, further systematic investigation is needed to determine whether the discrepancy reflects contextual sensitivity or methodological variation across studies.

## Conclusions

In summary, we show that modulation of pupillary hippus plausibly indicates motor-preparation. Hippus-based pupillometry therefore offers a scalable, noninvasive method for probing movement planning, with strong potential for supporting other measures of planning processes and enhancing e.g. brain-machine interface systems. Future progress will depend on combining hippus estimates of cognitive state and motor intention with other neural and behavioural modalities.

## Data availability statement

Data is available from the authors upon reasonable request via OSF: https://osf.io/ndfqa.

## References

[1] R. S. Johansson, G. Westling, A. Bäckström, and J. R. Flanagan, “Eye–Hand Coordination in Object Manipulation,” J. Neurosci., vol. 21, no. 17, pp. 6917–6932, Sept. 2001, doi: 10.1523/JNEUROSCI.21-17-06917.2001.

[2] “Motor Planning - Aaron L. Wong, Adrian M. Haith, John W. Krakauer, 2015.” Accessed: Dec. 11, 2025. [Online]. Available: https://journals.sagepub.com/doi/10.1177/1073858414541484

[3] S. Jainta, M. Vernet, Q. Yang, and Z. Kapoula, “The pupil reflects motor preparation for saccades - even before the eye starts to move,” Front. Hum. Neurosci., vol. 5, Sept. 2011, doi: 10.3389/fnhum.2011.00097.

[4] D. Voudouris, I. Schuetz, T. Schinke, and K. Fiehler, “Pupil dilation scales with movement distance of real but not of imagined reaching movements,” J. Neurophysiol., vol. 130, no. 1, pp. 104–116, July 2023, doi: 10.1152/jn.00024.2023.

[5] K. Fletcher, A. Neal, and G. Yeo, “The effect of motor task precision on pupil diameter,” Appl. Ergon., vol. 65, pp. 309–315, Nov. 2017, doi: 10.1016/j.apergo.2017.07.010.

[6] A. Yokoi and J. Weiler, “Pupil diameter tracked during motor adaptation in humans,” J. Neurophysiol., vol. 128, no. 5, pp. 1224–1243, Nov. 2022, doi: 10.1152/jn.00021.2022.

[7] A. Yokoi and J. Weiler, “Pupil diameter tracked during motor adaptation in humans,” J. Neurophysiol., vol. 128, no. 5, pp. 1224–1243, Nov. 2022, doi: 10.1152/jn.00021.2022.

[8] R. Johnston and M. A. Smith, “Brain-wide arousal signals are segregated from movement planning in the superior colliculus,” eLife, vol. 13, Oct. 2025, doi: 10.7554/eLife.99278.2.

[9] S. Mathot, “Pupillometry: Psychology, Physiology, and Function,” J. Cogn., vol. 1, no. 1, Feb. 2018, doi: 10.5334/joc.18.

[10] S. P. Marshall, “The Index of Cognitive Activity: measuring cognitive workload,” in Proceedings of the IEEE 7th Conference on Human Factors and Power Plants, Scottsdale, AZ, USA: IEEE, 2002, pp. 7-5-7–9. doi: 10.1109/HFPP.2002.1042860.

[11] A. T. Duchowski et al., “The Index of Pupillary Activity: Measuring Cognitive Load vis-à-vis Task Difficulty with Pupil Oscillation,” in Proceedings of the 2018 CHI Conference on Human Factors in Computing Systems, in CHI ‘18. New York, NY, USA: Association for Computing Machinery, Apr. 2018, pp. 1–13. doi: 10.1145/3173574.3173856.

[12] S. P. Marshall, “Whitepaper ICA,” US6090051A, July 18, 2000 Accessed: Apr. 10, 2025. [Online]. Available: https://patents.google.com/patent/US6090051A/en

[13] “Unity Real-Time Development Platform | 3D, 2D, V. & AR Engine,” Unity. Accessed: Apr. 25, 2025. [Online]. Available: https://unity.com

[14] “Hurricane VR - Physics Interaction Toolkit | Physics | Unity Asset Store.” Accessed: Jan. 04, 2025. [Online]. Available: https://assetstore.unity.com/packages/tools/physics/hurricane-vr-physics-interaction-toolkit-177300

[15] A. T. Duchowski, K. Krejtz, N. A. Gehrer, T. Bafna, and P. Bækgaard, “The Low/High Index of Pupillary Activity,” in Proceedings of the 2020 CHI Conference on Human Factors in Computing Systems, in CHI ‘20. New York, NY, USA: Association for Computing Machinery, Apr. 2020, pp. 1–12. doi: 10.1145/3313831.3376394.

[16] “jamovi - open statistical software for the desktop and cloud.” Accessed: Dec. 10, 2025. [Online]. Available: https://www.jamovi.org/

[17] A. E. Urai, A. Braun, and T. H. Donner, “Pupil-linked arousal is driven by decision uncertainty and alters serial choice bias,” Nat. Commun., vol. 8, no. 1, p. 14637, Mar. 2017, doi: 10.1038/ncomms14637.

[18] J. Beatty, “Phasic Not Tonic Pupillary Responses Vary With Auditory Vigilance Performance,” Psychophysiology, vol. 19, no. 2, pp. 167–172, 1982, doi: 10.1111/j.1469-8986.1982.tb02540.x.

[19] P. R. K. Turnbull, N. Irani, N. Lim, and J. R. Phillips, “Origins of Pupillary Hippus in the Autonomic Nervous System,” Invest. Ophthalmol. Vis. Sci., vol. 58, no. 1, pp. 197–203, Jan. 2017, doi: 10.1167/iovs.16-20785.

[20] S. Nobukawa et al., “Pupillometric Complexity and Symmetricity Follow Inverted-U Curves Against Baseline Diameter Due to Crossed Locus Coeruleus Projections to the Edinger-Westphal Nucleus,” Front. Physiol., vol. 12, Feb. 2021, doi: 10.3389/fphys.2021.614479.

[21] R. S. Larsen and J. Waters, “Neuromodulatory Correlates of Pupil Dilation,” Front. Neural Circuits, vol. 12, Mar. 2018, doi: 10.3389/fncir.2018.00021.

[22] A. Pomè, D. C. Burr, A. Capuozzo, and P. Binda, “Spontaneous pupillary oscillations increase during mindfulness meditation,” Curr. Biol., vol. 30, no. 18, pp. R1030–R1031, Sept. 2020, doi: 10.1016/j.cub.2020.07.064.

[23] S. R. Steinhauer, G. J. Siegle, R. Condray, and M. Pless, “Sympathetic and parasympathetic innervation of pupillary dilation during sustained processing,” Int. J. Psychophysiol., vol. 52, no. 1, pp. 77–86, Mar. 2004, doi: 10.1016/j.ijpsycho.2003.12.005.

[24] S. Nobukawa et al., “Pupillometric Complexity and Symmetricity Follow Inverted-U Curves Against Baseline Diameter Due to Crossed Locus Coeruleus Projections to the Edinger-Westphal Nucleus,” Front. Physiol., vol. 12, Feb. 2021, doi: 10.3389/fphys.2021.614479.

[25] S. Mathôt, “Pupillometry: Psychology, Physiology, and Function,” J. Cogn., vol. 1, no. 1, p. 16, Feb. 2018, doi: 10.5334/joc.18.

[26] R. Johnston and M. A. Smith, “Brain-wide arousal signals are segregated from movement planning in the superior colliculus,” bioRxiv, pp. 2024–04, 2025.

